# Serotonin induces DOWN states across the anesthetized mouse forebrain

**DOI:** 10.64898/2026.07.14.738517

**Authors:** Rosa Großmann, Jorge F. Mejias, International Brain Laboratory, Zachary F. Mainen, Guido T. Meijer

**Affiliations:** Max Planck Institute for Human Cognitive and Brain Science, Leipzig, Germany; Cognitive and Systems Neuroscience Group, Swammerdam Institute for Life Sciences, University of Amsterdam, Amsterdam, The Netherlands; Champalimaud Foundation, Lisbon, Portugal; Donders Centre for Neuroscience, Radboud University Nijmegen, The Netherlands

## Abstract

Serotonergic neurons in the dorsal raphe nucleus (DRN) project extensively throughout the forebrain, yet their influence on global network states remains controversial. Serotonin is implicated in regulating slow-oscillations, so-called UP and DOWN states, which occur during sleep and light anesthesia. While optogenetic fMRI suggests that serotonin (5-HT) suppresses brain-wide activity, classical electrical stimulation studies report the induction of cortical UP states. To resolve this discrepancy, we combined optogenetic 5-HT stimulation with large-scale Neuropixel recordings across the forebrain of lightly anesthetized mice. We demonstrate that selective 5-HT release consistently induces DOWN states across the cortex and striatum, while leaving midbrain dynamics largely unaffected. A multi-area computational model constrained by biologically plausible connectivity, indicates that in certain regions the transitions arises from network-level synchronization rather than direct DRN input.

## Introduction

Serotonin (5-HT) is a major central neuromodulator that orchestrates a vast array of brain states and cognitive functions [1]. Among its targets are slow oscillations; rhythmically alternating states of high neural activity (UP states) and quiescence (DOWN states). Slow oscillations appear during anesthesia and slow-wave (non-REM) sleep and are important for learning, memory consolidation, and plasticity [2–5], they are considered to be the default emergent activity of neural networks [6].

The current literature offers a fractured landscape of serotonergic effects on slow oscillations. Evidence suggests that tonic increases in 5-HT, such as those induced by MDMA or bath application in slices, can suppress slow oscillations entirely [7]. Furthermore, short transient pulses of 5-HT, by optogenetically activating serotonergic neurons in the DRN, suppress neural activity across the anesthetized brain, as measured with fMRI [8, 9]. Electrophysiological recordings in the frontal cortex similarly revealed a serotonin-induced suppression of neural activity [9]. However, opposing results have occasionally been reported, such as the induction of UP states following electrical stimulation of the dorsal raphe nucleus (DRN) [10]. Serotonin clearly affects slow oscillations, but its precise impact of serotonin on UP-DOWN dynamics remains debated.

Resolving the ambiguity of serotonin’s role requires an approach that matches its global reach while measuring neural activity with high temporal precision. While fMRI has successfully mapped brain-wide serotonergic suppression under anesthesia [8, 9], it lacks the temporal resolution to resolve UP and DOWN states. Conversely, the electrophysiological studies capable of capturing these fast dynamics have remained restricted to localized cortical regions, leaving the question of brain-wide coordination unanswered [9]. Here, we bridge this gap by employing Neuropixel probes to record large-scale single-cell spiking activity across the cortex, amygdala, striatum, hippocampus, thalamus, and midbrain during optogenetic activation of DRN serotonergic neurons.

We found that 5-HT stimulation rapidly induced a DOWN state in cortex, amygdala, striatum, hippocampus, thalamus, but not the midbrain. Computational modelling of UP-DOWN dynamics across all regions, linked together using realistic inter-area connectivity, could replicate the experimental data. The model revealed that cortical regions which do not receive direct DRN input—like the visual cortex—were still transitioned into a DOWN state through network-level interactions.

## Results

### Serotonin stimulation suppresses spiking activity across the mouse forebrain under anesthesia

We performed acute Neuropixel recordings across the mouse brain while the animal was under light isoflurane anesthesia. Serotonin was stimulated by activating serotonergic neurons in the DRN optogenetically during Neuropixel recordings in the forebrain (Fig. 1a,b). Over four consecutive recording days, seven Neuropixel insertions were made targeting frontal cortex, visual cortex, piriform cortex, amygdala, striatum, hippocampus, thalamus, and the midbrain. (Fig. 1c) Serial-sectioning two-photon microscopy was used to reconstruct the entire brain volume post-mortem. The reconstructed brain volume was used to determine channelrhodopsin expression in the DRN and reconstruct the Neuropixel insertion trajectories across the brain (Fig. S1). Out of seven mice, five showed good expression of channelrhodopsin in the DRN and were included in subsequent analysis, the two lacking expression, most likely due to mistargeted viral injections, were used as controls.

**Figure 1.**
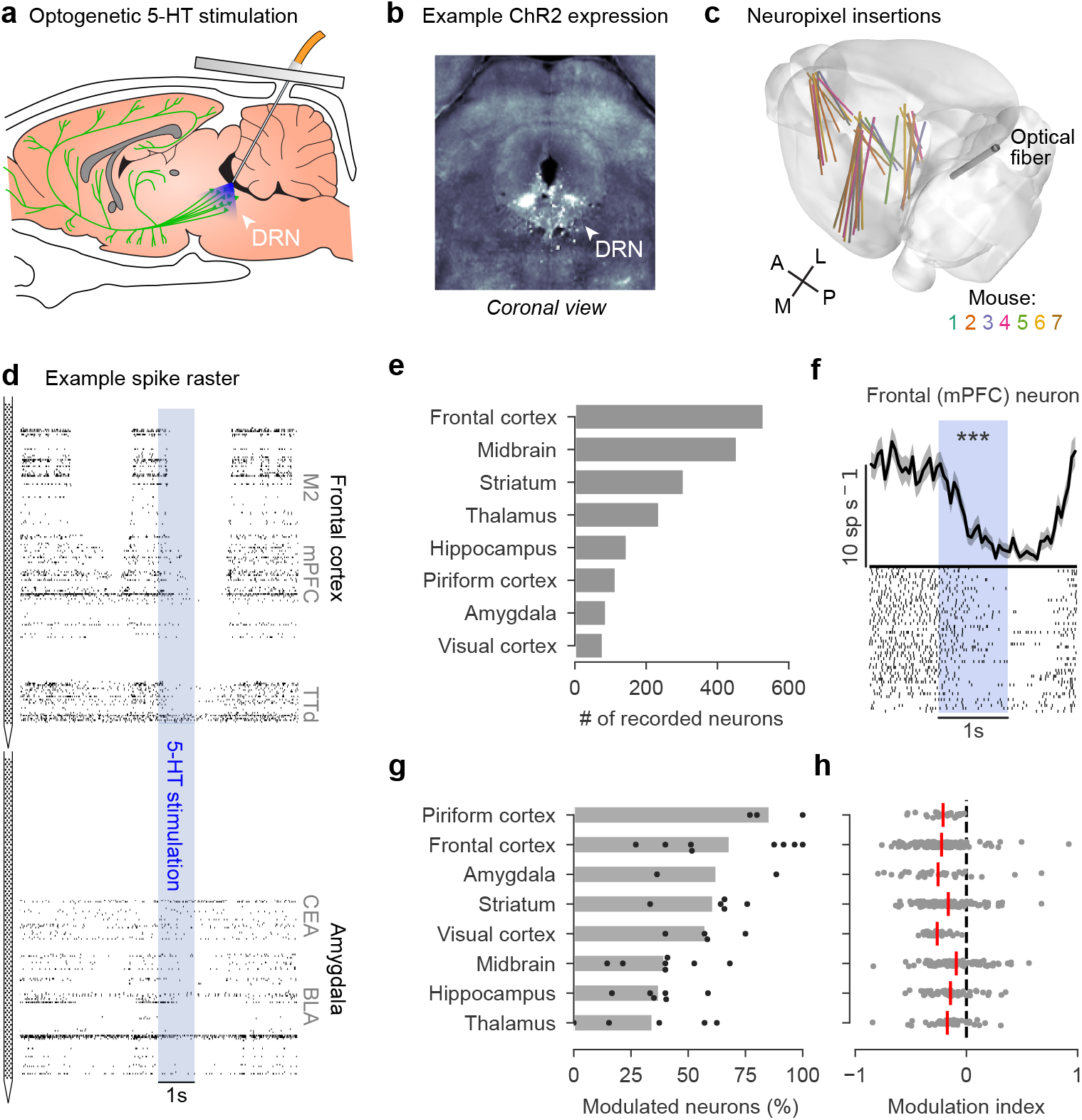
Serotonin stimulation inhibits spiking activity across the brain. **(a)** Saggital cartoon drawing of serotonergic neurons in the DRN projecting throughout the brain (green). An optical fiber is positioned directly above the DRN through the cerebellum through which the 5-HT neurons can be activated with blue light. **(b)** A coronal histological section of the DRN showing expression of channelrhodopsin (ChR2). **(c)** Neuropixel insertion trajectories, aligned to the Allen Brain reference atlas, across all seven mice. Seven insertions were made per mouse over a period of four days, typically two at the same time. Color of each insertion indicates mouse identity. **(d)** An example spike raster plot of 10s of spiking activity recorded on two Neuropixel probes simultaneously. Recorded brain regions are indicated on the right of the raster at their approximate depths. The time of the optogenetic 5-HT stimulation is indicated by the blue bar (25 Hz, 10 ms pulses). **(e)** The total number of recorded neurons across all recording sessions split out by brain region. **(f)** Peri-stimulus time histogram of an example neuron from the frontal cortex. The bottom shows the spike raster in which each row is a repetition of 1s 25 Hz stimulation. The top shows the mean spiking rate binned in 50 ms bins with a 25 ms Gaussian smoothing kernel. *** *p <* 0.001, ZETA-test. **(g)** Percentage of significantly modulated neurons (ZETA-test, *p <* 0.05) per brain region, calculated per recording session. Black dots are sessions, grey bars means over sessions. **(h)** Modulation index per neuron, negative values indicate the neuron is inhibited by 5-HT and positive values indicate excitation. Each grey dot is a single neuron, vertical red bars are the mean over neurons per brain region.

Cortical UP-DOWN states were clearly visible under isoflurane anesthesia as periods of 1-2s of neural activity interspersed with periods of silence in cortex, striatum, and thalamus (Fig. 1d). Based on visual inspection of the spike raster plots, 5-HT stimulation appeared to coincide with a DOWN state. To investigate this, first we asked what the effect of serotonin stimulation was on the activity of single neurons, which were recorded in large numbers across the targeted brain regions (Fig. 1e). Peri-stimulus time histograms of single neurons, conditioned on the start of the 5-HT stimulation, consistently showed that single neuron spiking activity was decreased (example neuron in Fig. 1f). In all recorded brain regions, we found a large fraction of neurons that were significantly modulated by serotonin stimulation (30-60%; ZETA-test [11], *p <* 0.05; Fig. 1g). A large majority of single neurons were suppressed by 5-HT stimulation as quantified by their modulation index, which was predominantly *<* 0. The mean modulation index over neurons per brain region was negative indicating that all recorded regions were suppressed by serotonin stimulation (Fig. 1h).

### Serotonin stimulation induces a DOWN state

To investigate whether serotonin stimulation induced a DOWN state we quantified UP and DOWN states with a two-state Hidden Markov Model (HMM) which was fit to spike rates conditioned to the start of the 5-HT stimulation (Fig. 2a,b). We found that serotonin stimulation strongly and rapidly induced a DOWN state in the frontal cortex, this effect scaled with the stimulation frequency of the optogenetic stimulation (Fig. 2c). Overall, brain regions which naturally show bi-stable dynamics under anesthesia—cortex, striatum, and thalamus—were transitioned into a DOWN state when stimulating 5-HT neurons in the DRN (Fig. 2d). Besides these, also brain regions which do not show endogenous UP-DOWN state dynamics—amygdala and hippocampus—showed a 5-HT induced increase in DOWN state occurrence. This DOWN state likely reflects a suppression of neural firing but does not constitute the induction of a naturally occurring DOWN state, since these regions do not show them normally. The midbrain does not show UP-DOWN state dynamics normally and serotonin stimulation also did not induce a DOWN state as it did in all the other brain regions. Besides the induction of a DOWN state, serotonin stimulation also induces a rebound UP state in thalamus, frontal cortex, and to a lesser extend in the midbrain. The rebound UP state in the midbrain is particularly interesting since no DOWN state was induced there. Importantly, these results could not be explained by neural responses to the optogenetic light delivery alone (Fig. S2).

**Figure 2.**
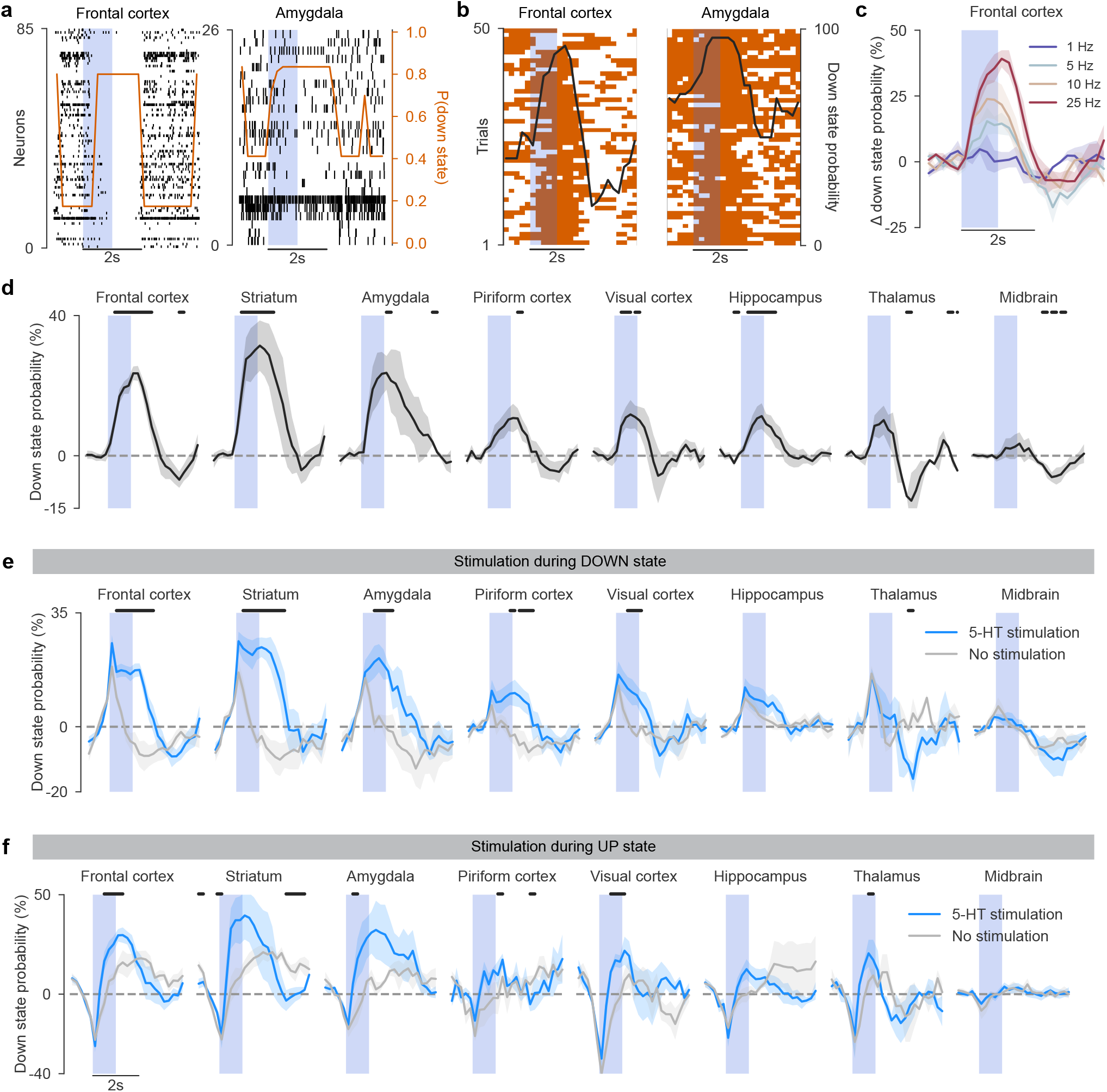
Serotonin induces a DOWN state under anesthesia. **(a)** An example peri-stimulus raster plots of spike times (black ticks, each row is a neuron), aligned to the onset of 5-HT stimulation (blue vertical bar). The orange line indicates the posterior probability of a DOWN state as determined by a two-state HMM. On the left an example from the frontal cortex is shown and on the right an example from the amygdala. **(b)** A peri-stimulus heatmap over trials of when the HMM determined there most likely was a DOWN state (orange). The black line is the mean posterior probability of a DOWN state over trials per time bin. **(c)** A peri-stimulus plot of the change in DOWN state probability compared to baseline (−1 to 0s) averaged over mice (n = 5, mean ± sem). Plotted separately, indicated by color, are different stimulation frequencies. **(d)** Peri-stimulus plots for all recorded high-level brain regions of the change in DOWN state probability over baseline for 25 Hz serotonin stimulation. Black horizontal bars above the plot indicate periods of statistical significance (t-test versus 0, *p <* 0.05). **(e)** Peri-stimulus plots of the change in DOWN state probability over baseline, subselected for optogenetic stimulation bouts that started during a naturally occurring DOWN state (5-HT stimulation; blue line). The grey line shows the DOWN state probability over baseline for randomly jittered times, subselected DOWN states. Black horizontal bars show the periods of statistical difference between 5-HT stimulation and no stimulation (paired t-test, *p <* 0.05). **(f)** Same as (e) but for stimulation times subselected to occur during an ongoing UP state.

The observed increase in DOWN state probability could result from a lengthening of an ongoing DOWN state, an abrupt transition to a DOWN state from an ongoing UP state, or both. To disentangle these possibilities we split the stimulation bouts into stimulation which occurred during an ongoing DOWN or UP state. When stimulation occurred during an ongoing DOWN state, this DOWN state became longer (Fig. 2e). Conversely, when 5-HT stimulation occurred during an ongoing UP state, the UP state rapidly ended, and a DOWN state started (Fig. 2f). This showed that serotonin stimulation resulted in both longer ongoing DOWN states and rapid transitions to DOWN states from ongoing UP states.

We next investigated the contribution of different serotonergic receptor subtypes to the magnitude of the 5-HT induced suppression of neural activity. It was shown *in vitro* that blocking 5-HT1a and 5-HT3 receptors did not impact serotonergic suppression of slow-oscillations whereas blocking 5-HT2a receptors did [7]. Also correlative results, obtained with fMRI, showed a link between 5-HT2a receptor expression and the strength of serotonin-induced suppression of neural activity at the brain region level [9]. Strikingly, there was no correlation between the strength the anatomical projection of the DRN to a given area and the magnitude by which it was suppressed by optogenetic stimulation of serotonergic neurons in the DRN [9]. Here we investigated this by obtaining the DRN projection strength and receptor expression profiles for various serotonergic receptors from the Allen Brain Atlas and correlated them with the magnitude of the suppression of spiking activity elicited by serotonin stimulation at a region-by-region basis. We found no significant correlation with the DRN projection strength (Fig S3a). Out of all the serotonergic receptors expression profiles we were able to obtain from the Allen Brain Atlas, we found significant correlations with 5-HT1f and 5-HT2a expression, such that higher levels of receptor expression were associated with stronger suppression of neural activity in that region (Fig S3b-d). In short, we replicated previous results obtained in slice and with fMRI and found an additional link to 5-HT1f receptor expression.

### Computational modelling of UP-DOWN state dynamics in a bistable regime

We aimed to shed further light on the mechanisms underlying serotonergic suppression of neural activity and the induction of DOWN states across the brain. To this end we implemented a dynamic bistable firing rate model to investigate the mechanism by which serotonin stimulation results in the induction of DOWN states. The model simulates a simplified network of the mouse brain, following biological realistic inter-area connectivity [12, 13]. Each brain area was modelled using a local excitatory-inhibitory (EI) circuit consisting of one E, one I population, and a mechanism controlling for adaptation (A) of the excitatory population [14] (Fig. 3a). To introduce biologically plausible connectivity between the modelled brain regions, the projection strength between them was pulled from the Mouse Connectivity Atlas of the Allen Brain Atlas [12, 13], together with the projection strength of the dorsal raphe nucleus to these regions (Fig 3b). The model was able to reproduce slow-wave oscillations which mimicked closely the UP and DOWN states observed in the neural data (Fig. 3c). The frequency of modelled UP and DOWN states was around 0.5-1 Hz which closely matched the frequency observed in the brain.

**Figure 3.**
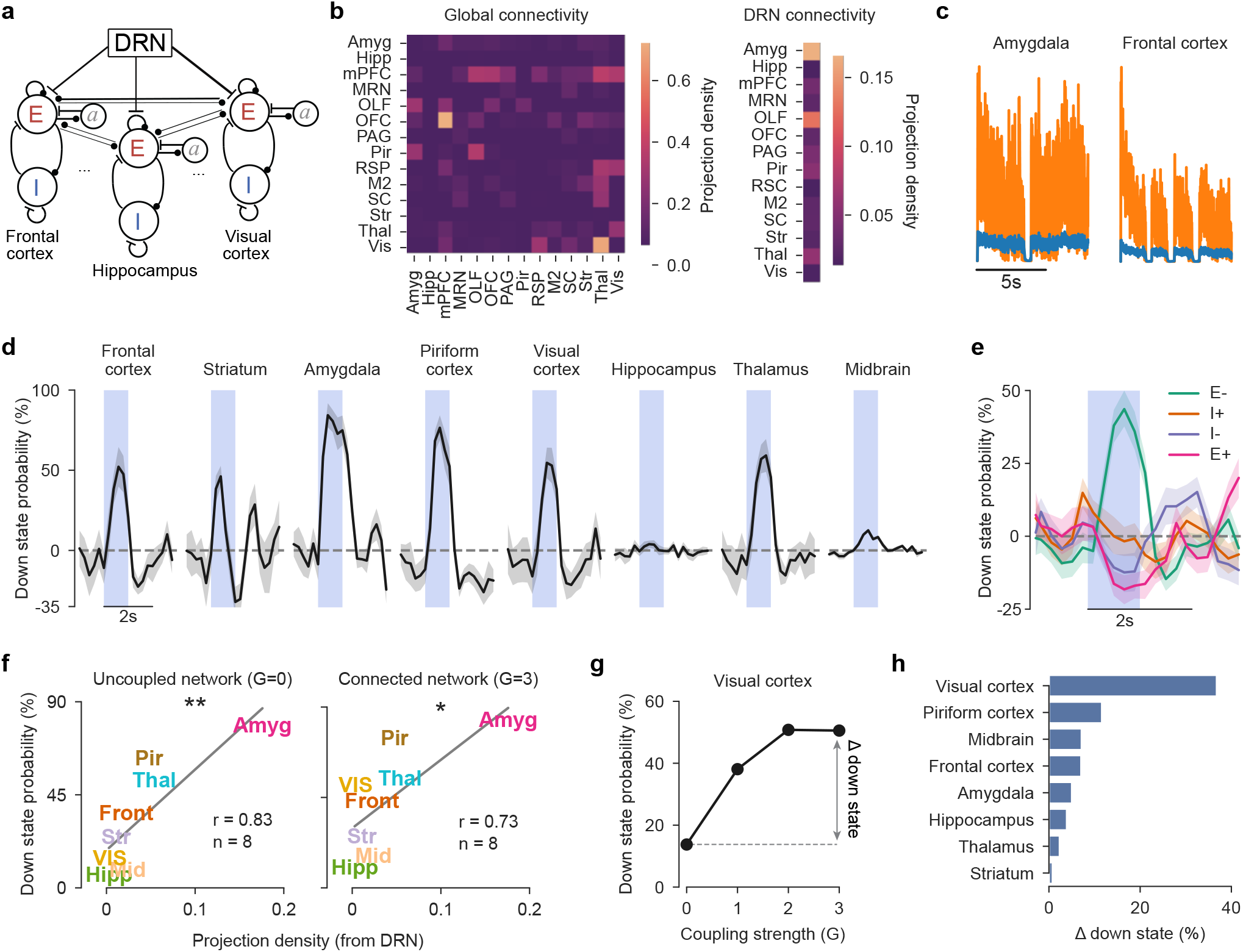
Simulation of UP-DOWN state dynamics by a bistable firing rate model. **(a)** A global circuit model of interconnected brain regions which receive input from the dorsal raphe nucleus (DRN). **(b)** Connectivity matrix for the global circuit and DRN projections. **(c)** The modelled activity recapitulated UP-DOWN state dynamics in brain regions which also showed these dynamics in the brain; e.g. the frontal cortex. **(d)** Modelled serotonin stimulation induced a DOWN state in several brain regions. The change in DOWN state probability versus baseline (−1 to 0s) is plotted per brain region. **(e)** DOWN states were induced by inhibiting the excitatory population (E−), but not by exciting the inhibitory population (I+), inhibiting the inhibitory population (I−), or exciting the excitatory population (E+). The change in DOWN state probability is plotted as in panel d, averaged over regions. **(f)** In an uncoupled network (coupling strength *G*=0, left panel), the DRN projection density to a given brain region determines the strength of the effect of the modulation by serotonin. Brain regions (n=8) are colored words. Amyg: amygdala, Pir: piriform cortex, Thal: thalamus, Front: frontal cortex, Str: striatum, VIS: visual cortex, Mid: midbrain, Hipp: hippocampus. ** *p*=0.01, r=0.83, Pearson correlation. In a network with strong realistic connectivity between brain regions (*G*=3, right panel) the correlation was less strong. * *p*=0.04, r=0.73, Pearson correlation. A leave-one-out analysis showed that the correlation in the uncoupled network (*G*=0) survived removal of any single region, whereas the correlation in the coupled network (*G*=3) did not survive this analysis and relied primarily on the amygdala. **(g)** The probability of a DOWN state, as defined by the maximum during 5-HT stimulation (0-1s), plotted for increasing values of inter-area coupling strength *G* in the network. Here only visual cortex is plotted. The effect of coupling strength on DOWN state probability is defined as the absolute difference between *G* = 0 and *G* = 3. **(h)** The Δ down state, as defined in panel **(g)**, per brain region.

Serotonin stimulation was modelled by targeting the E population of each brain region with an inhibitory current (E−) proportional to the projection strength from the DRN to that region. This implementation was chosen because inhibiting E increases the firing threshold, making it likely to induce a transition from the UP to the DOWN state. To investigate the effect on UP-DOWN state dynamics, a two-state HMM was fitted to the model output, as for the recorded data. Serotonin stimulation induced a DOWN state across brain regions in a qualitatively similar manner to the neural data (Fig. 3d). One notable exception was the hippocampus, which showed weak DOWN state induction in the neural data but none at all in the model.

Three alternative implementations of serotonin stimulation are also plausible: an excitatory current targeting the I population (I+), which should intuitively produce a similar outcome to E-; an excitatory current targeting E (E+); or an inhibitory current targeting I (I−). Running the model with all four scenarios revealed that only E-resulted in a transition to a DOWN state (Fig. 3e). Contrary to our expectation, I+ did not induce a DOWN state. Inspecting the nullclines of the excitatory and inhibitory populations of one isolated area revealed one possible explanation: at equal input strength, the I-condition leads to a single stable fixed point (the DOWN state), whereas the I+ condition remains bistable, leaving the option to stay in an UP state open (Fig. S4). Forcing a DOWN state via I+ might therefore be theoretically possible, but more difficult to achieve.

We next investigated the importance of the inter-areal coupling strength in the model, given its importance in stable signal propagation across brain networks ([15]), by varying the coupling strength *G*. In a completely uncoupled network (*G* = 0), the probability of a DOWN state in each region during simulated 5-HT stimulation was strongly correlated with the projection from the DRN (Pearson correlation, *r*=0.83, *p*=0.01; Fig. 3f, left). Since the correlation is only based on eight regions, we performed a leave-one-out analysis, showing that the correlation is robust and does not rely on a specific region (Fig. S5a). In the case of high global coupling a significant correlation remained when including eight areas (*r*=0.73, *p*=0.04; Fig. 3f, right), but the leave-one-out analysis revealed that this correlation was not robust and largely driven by the amygdala (Fig. S5b).

We observed in Fig. 3f that some regions showed a large change in the probability of a DOWN state between the uncoupled and connected model whereas other regions did not show any difference. This was quantified by taking the Δ down state between the *G* = 0 and *G* = 3 models for each brain region (Fig. 3g). This revealed that the visual cortex showed the largest change in DOWN state probability compared to the other brain regions by far (Fig. 3h). Notably, it was not the case that any region receiving weak DRN input showed a large change in down state probability since the Δ down state was not correlated with DRN input strength (Fig. S3a; Fig. S5c). This means that the 5-HT induced DOWN state in visual cortex is not due to a direct projection from the DRN but instead because the visual cortex starts following the UP-DOWN state dynamics of the rest of the cortex through inter-area communication.

## Discussion

Here we showed that selective activation of serotonergic neurons in the DRN under light isoflurane anesthesia resulted in an inhibition of neural activity across the mouse brain and a rapid induction of a DOWN state across large parts of the mouse forebrain. Doing so bridged the gap between whole-brain fMRI studies and localized electrophysiological results from single brain regions by performing large-scale and widespread Neuropixel recordings. Computational modelling further revealed that cortical regions which lack direct DRN input—like the visual cortex—are still transitioned into a DOWN state due to network-level interactions.

Slow oscillations, or UP-DOWN states, are observed in the cortex, thalamus and striatum during non-REM sleep. Contrary to REM sleep, during which serotonin is tonically low, serotonin levels in cortex show phasic peaks during non-REM sleep [16]. This suggests that, during non-REM sleep, serotonin serves as a state switch between periods of UP-DOWN state dynamics and periods of overall suppression of neural activity. Whether isoflurane-induced UP-DOWN state dynamics are mechanistically equivalent to those observed during natural slow-wave sleep is debatable. However, a direct comparison of UP-DOWN state characteristics between non-REM sleep and light isoflurane anesthesia showed that they are fairly similar [17].

In conflict with our results, electrical stimulation of the DRN was shown to induce an UP state in frontal cortex in rats under chloral hydrate anesthesia [10]. The most likely explanation for the contradictory results is that electrical stimulation of the DRN is non-selective and activates glutamatergic as well as serotonergic neurons. The glutamatergic drive is likely much stronger compared to the serotonergic one and dominates the down-stream effect resulting in the induction of an UP state instead of a DOWN state. Another possibility, albeit less likely, is that the study of [10] was performed in rats whereas the other studies, including this one, were performed in mice. To our knowledge, however, there is no reason for the effect of serotonin to be opposite in rats versus mice.

The amygdala and hippocampus do not normally show UP-DOWN state dynamics, however, they did show increased DOWN state probability in response to 5-HT stimulation. This is likely not the system transitioning into a DOWN state—since they do not naturally occur—but simply a suppression of neural activity which is picked up by the two-state HMM as a DOWN state. This suggests that the induced DOWN states in the amygdala and hippocampus are qualitatively different from those that are induced in regions which do naturally exhibit UP-DOWN dynamics, such as the cortex, thalamus, and striatum. Follow-up experiments are necessary to show a mechanistic difference between the 5-HT induced DOWN states in regions which are naturally bi-stable and those which are not.

A limitation of our modelling approach was that the inter-area connectivity introduced to the model was open-loop; the DRN only provided input to downstream target regions, these regions did not project back to the DRN. We acknowledge this limitation but do not expect qualitative differences in the results for more realistic closed-loop architectures. One reason we do not expect this is that 5-HT input from the DRN shuts down its target regions, effectively preventing them from providing top-down feedback to the DRN.

Besides serotonin, some serotonergic neurons in the DRN co-release glutamate. Although the percentage is relatively low among serotonergic neurons projecting to subcortical areas (~ 15%), it is much higher in serotonergic neurons projecting to cortex (~ 60%; [18]). The co-release of glutamate should result in excitation of downstream neuronal populations, which is something we putatively observed in the awake condition [19]. During light anesthesia, however, the excitatory glutamatergic drive is masked by the dominant suppressive effect of serotonin release. Targeted pharmacological studies are required to determine exactly which biophysical mechanism causes the strong inhibitory drive under anesthesia. Our correlative results suggest that the 5-HT1f and 5-HT2a receptors are involved.

A surprising finding was that in the electrophysiological data there was no correlation between the strength of the 5-HT induced suppression and the projection strength of the DRN to that region (Fig. S3a). In the awake condition we found the same result [19]. Our computational model provided a possible explanation; when increasing the inter-area connectivity in the model the correlation with DRN projection strength went down (Fig. 3f). In the computational model, however, the correlation with DRN input never completely disappeared. This was likely due to the fact that the model was a simplified version of reality which lacked physiological detail, such as serotonergic receptor expression patterns and volume transmission of serotonin across region boundaries. Regardless, the model showed that a subset of brain regions, lacking direct DRN input, are still transitioned into a DOWN state through network-level synchronization of neural activity.

## Data and code availability

All data are openly available through the Open Neurophysiology Environment (ONE) protocol (https://int-brain-lab.github.io/ONE/). All the code used to generate the figures in this manuscript are on Github: https://github.com/rosagross/SerotoninAnaesthesiaModel/.

## Acknowledgments

We thank the staff of the International Brain Laboratory collaboration for their technical support. This work was supported by the Wellcome Trust (209558 and 216324) and the Simons Foundation.

## Material and methods

### Animals

Seven SERT-Cre C57Bl/6 mice (four males and three females) were used for the purpose of this study. All experimental procedures were approved by the Portuguese Veterinary General Board.

### Surgical procedures

Mice underwent two surgeries: (1) injection of the channelrhodopsin viral vector in the dorsal raphe nucleus and implantation of the optical fiber & headbar, and (2) the creation of three small craniotomies in the skull. Both these surgical procedures were performed as previously described in [19]. A minimum of two weeks was allowed for the virus to express and the mouse to recover after the first surgery. The second surgery was performed ~ 12 hours preceding the first Neuropixel recording.

### Neuropixel recordings

Acute anesthetized Neuropixel recordings were performed on four consecutive days. Mice were anesthetized with an induction dose of 3% isoflurane after which they were headfixed in the recording rig (hardware described in detail in [19]). A maintenance level of isoflurane of 0.8-1.5% was administered through a mask which was placed over the nose of the mouse. The tip of each Neuropixel probe was lowered in drop of Di-I with the micromanipulator prior to insertion. One or two Neuropixel probes, depending on the recording day, were inserted in the following coordinates:

| Session | ML ($\mu\text{m}$) | AP ($\mu\text{m}$) | Depth ($\mu\text{m}$) | Angle ($^{\circ}$) | Directionality |
| --- | --- | --- | --- | --- | --- |
| 1 | -1090 | 2000 | 3800 | 15 | medial |
| 1 | -2184 | -2000 | 6800 | 10 | lateral |
| 2 | -946 | -3300 | 3800 | 10 | medial |
| 2 | -1227 | 2000 | 4590 | 10 | lateral |
| 3 | -2240 | -2000 | 4000 | 15 | medial |
| 3 | -850 | -3460 | 5270 | 10 | lateral |
| 4 | -2250 | -1800 | 5500 | 17 | lateral |

Throughout the recording session, the isoflurane level was calibrated such that the mouse would not wake up but UP and DOWN states were visible in the electrophysiological recordings. Once mice were stable under light anesthesia, serotonergic neurons in the dorsal raphe nucleus were optogenetically stimulated with 1s pulse trains of blue light (465 nm) administered through the optical fiber. The light power was 5 mW, as measured at the tip of the optical fiber. Light pulses were 10 ms long and with a frequency of 1, 5, 10, or 25 Hz. Different frequencies were administered in a randomized order and repeated each 50 times. The inter-stimulus interval was drawn from a truncated exponential distribution with a factor of 6s, a minimum of 5s, and a maximum of 15s. The whole stimulation protocol lasted around 30 minutes, during this time the body temperature of the mice was kept from falling using hand warmer packages (electrical heating pads produced large artifacts in the electrophysiological recordings). After the protocol, the Neuropixels were retracted, the skull covered, and the animal was placed in a cage positioned on a heating pad to wake up.

### Processing of electrophysiological data

Pre-processing and spike sorting of raw electrophysiological data was performed with the fully automated pipeline developed by the International Brain Laboratory (IBL) [20]. We only included neurons that passed all the IBL neuron-level quality control criteria. Additionally we excluded neurons with firing rates *<* 0.1 spks/s over the whole recording.

### Modulation index

For each neuron a modulation index was calculated which quantifies the strength and directionality of the modulatory effect of serotonin. A 500 ms baseline window was defined directly before stimulus onset, and a 500 ms stimulus window was defined at 300 - 800 ms after stimulus onset. The modulation index *MI* was defined as the area under the ROC curve *auROC* between the spike counts in the baseline window and the stimulus window, which was scaled between −1 and 1 according to Equation 1.

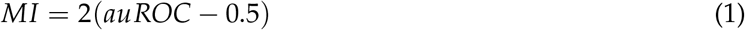

### Firing rate model

We implemented a bistable firing rate model, consisting of multiple local circuits which were all-to-all connected to form a global network. The local circuits represented the relevant brain areas from the experimental recordings. Serotonin stimulation was modelled by inducing a current to all regions, originating from the DRN. The connectivity was realistically modelled by using connectivity data from the Allen Brain Atlas [12, 13] which defined the connectivity between the regions in the model as well as the projection density from the DRN to the different regions.

The local circuit model was based on the implementation by Jercog et al. [14], including one excitatory population *E* and one inhibitory population *I*. It included an adaptation module *a*, simulating synaptic fatigue in pyramidal cells, as well as background noise which was introduced to the E and I population. Both populations projected to each other, as well as to themselves through self-coupling. The following equations describe the change of the populations *E* and *I* firing rate *r* (see Equation 2 and 3), as well as the strength of adaptation *a* (see Equation 4) over time, where *τ*_*E*_, *τ*_*I*_ and *τ*_*a*_ are time constants:

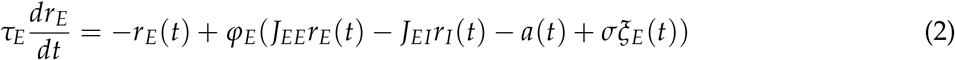

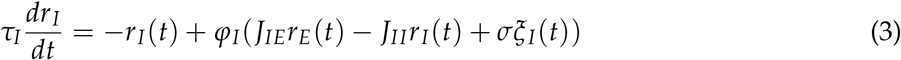

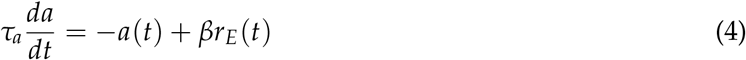

The frequency of UP and DOWN state changes was defined by 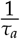, so the original value of *τ*_*a*_ = 500*ms* generated states in the frequency of around 2Hz. Analyses of the experimental data in the mouse revealed that the frequency of state change is 0.5-1Hz, therefore a value of *τ*_*a*_ = 1500*ms* was chosen and the other time constants were scaled accordingly, so that the final implementation yielded *τ*_*E*_ = 30*ms* and *τ*_*I*_ = 6*ms*. The variable *J* defined the local coupling strength for the E and I population, which was *J*_*EE*_ = 5, *J*_*EI*_ = 1, *J*_*IE*_ = 10, and *J*_*II*_ = 0.5 (the first letter indicating the target population). To model the noise of the system, which was introduced as a background noise to both populations, two Ornstein-Uhlenbeck processes were implemented, with a mean of zero and a standard deviation of *σ* = 3.5. The functions *φ*_*E*_ and *φ*_*I*_ were the transfer functions, which determined the reaction of the population to incoming currents, through self coupling, projections coming from other populations, or/and background noise. The equation is a threshold-linear function, with a threshold that defines the start of the linear part of the equation. If the input did not reach the threshold, the activation of the regarding population was set to zero.

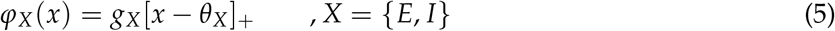

The threshold that has to be reached until the population reacts to incoming input is described by *θ*. For all brain regions we kept the original *θ*_*I*_ = 25 as threshold for the inhibitory population and used a region specific threshold for the excitatory population (see below). In case the input reached the threshold level, the slope *g* determined the output. *g* controlled how sensitive the population reacts to input; a higher *g* will lead to a stronger output than a lower *g*. The slope *g* was fine-tuned for each brain region so that the average firing rate of the E and I population is slightly below the value measured in the mouse brain, since inter-area connectivity further increases the baseline firing rate. UP and DOWN state characteristics were also matched to the experimental data on a region-by-region level; some brain regions, like the cortex, showed strong UP-DOWN dynamics *in vivo* whereas other regions, like the midbrain, did not. To mimic this in the model, the adaptation strength *β* and the threshold of the E population *θ*_*E*_ were fine-tuned per brain region. For regions which showed only UP states we chose *β* = 1 and *θ*_*E*_ = *−* 1, regions with infrequent UP-DOWN dynamics were modelled at *β* = 3 and *θ*_*E*_ = 0, and regions with frequent UP and DOWN states at *β* = 6 and *θ*_*E*_ = *−* 1. Note that the parameter fine-tuning is a model fitting procedure rather than a validation of the experimental results. The models contribution lies in testing the relevance of interarea coupling in the serotonin effect.

### Modelling inter-area connectivity

Realistic connectivity was introduced to the model using data from the Mouse Connectivity Atlas by the Allen Brain Institute [12, 13]. Brain regions which were targeted by the Neuropixel recordings were defined as regions as interest. For each region, we obtained the projection density values to all other regions. Results from all experiments per region were averaged. Since some experiments did not show any projection values, a threshold of a projection density of 0.005 was applied for the experiment to be included. As a next step, the obtained values of the fine grained areas were grouped together into higher level brain areas that each represent one component of the network (Fig. 3b). Besides the connectivity values between the areas within the network, the anatomical projections originating from the DRN were set according to tracing data from the Allen Brain Institute (Fig. 3b). The connectivity data was introduced to the global network model, by taking the projection density values *W*^*xy*^ (projection to area *x* from area *y*) and multiplying them with the coupling strength *G* between the inter-area projections. The input into population E of area *x* can therefore be defined as 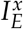 from projections that originate from all other areas y (see Equation 6).

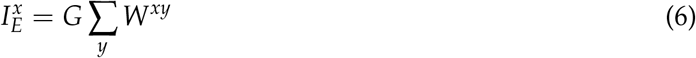

### Modelling 5-HT stimulation

The modulatory effect of serotonin stimulation was modelled by a current from the DRN projecting to the local circuits in the model. Similar to the connectivity matrix for the inter-area long-range projections, the projections of the DRN are weighted depending on the projection density. Considering that the DRN projection densities to all areas *x* are stored in a vector **D** the input current *I*_*DRN*_ is defined as the following:

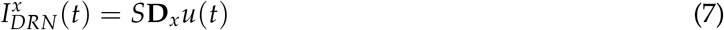

The temporal dynamics of the release of serotonin in the model was matched to the temporal profile of the modulatory effect of 5-HT on neural activity (data not shown). The trajectory of 5-HT modulation peaked at ~ 0.5 seconds and decayed back to baseline by ~ 3 seconds. The time course of this trajectory, here defined as *u*(*t*), was multiplied with the parameter for serotonin stimulation strength *S*. The stimulation was given for 1 second, with an inter-stimulus interval similar to the experimental protocol. The effect of serotonin stimulation on the system was initially modelled by introducing an inhibitory current to the E population (subtracting *I*_*DRN*_ from *r*_*E*_), this was later extended to other mechanisms.

### Hidden Markov Model (HMM)

In both the electrophysiological and simulated data, UP and DOWN states were quantified by fitting an HMM using the ssm package [21]. In the case of the electrophysiological data, spike times were binned in 200 ms bins in seven second windows (2s pre, 5s post) surrounding 5-HT stimulation onsets. An HMM with two states and Poisson observations was fit to the binned spike rates. In the case of the model, an HMM with two states and Gaussian observations was fit to the modelled firing rate time series of the I population. The state which showed the highest firing rate was determined to be the UP state. The output of the HMM was an inferred UP or DOWN state per time point and a corresponding posterior likelihood.

## Supplementary figures

**Figure S1.**
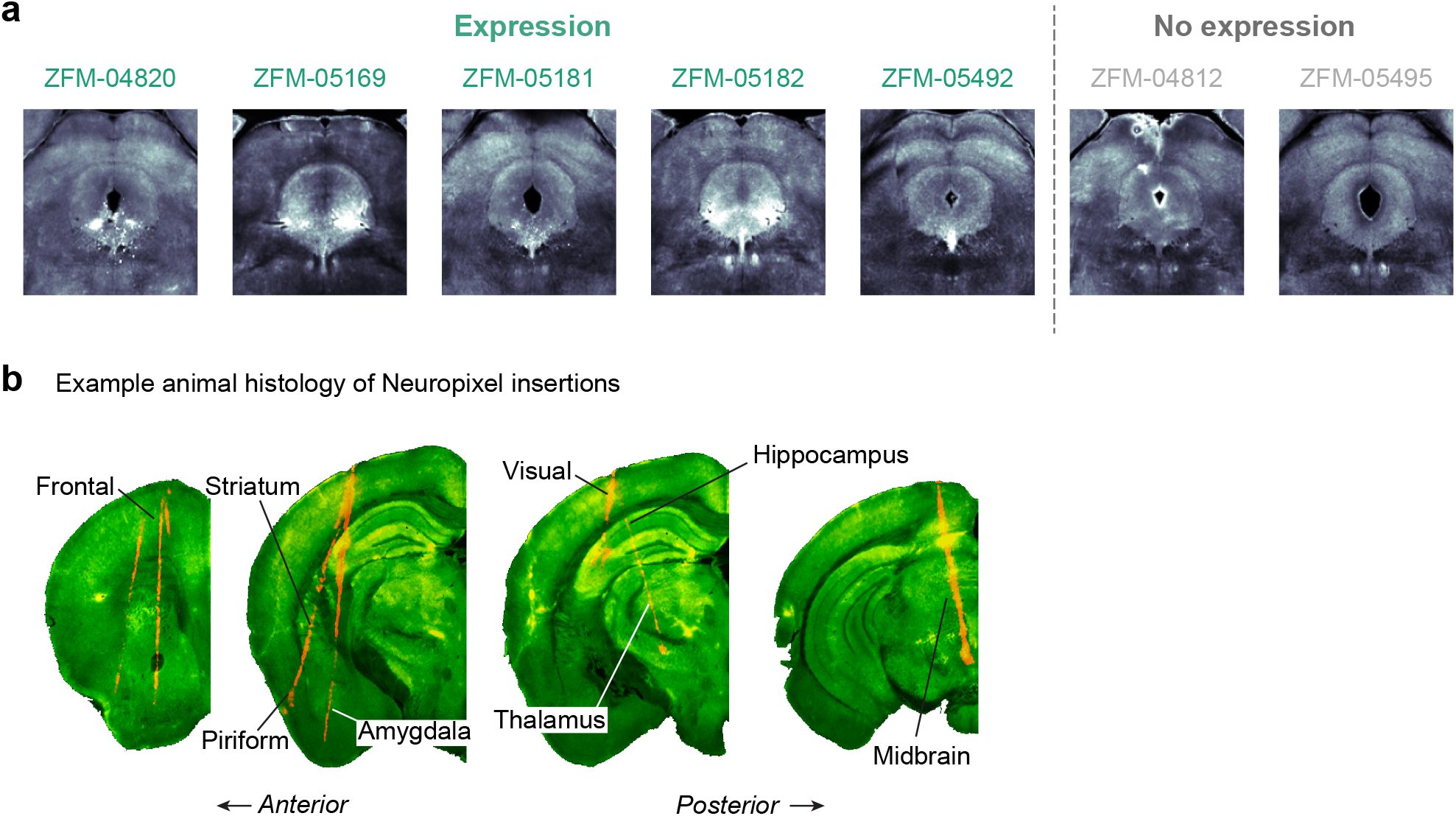
Histology. **(a)** Coronal slices of channelrhodopsin expression in the DRN for all seven SERT-cre mice. The first five mice, labeled in green, show expression of channelrhodopsin in the DRN as indicated by fluorescence directly underneath the ventricle. The last two mice, labeled in gray, do not show any expression due to a failed virus injection. **(b)** Fluorescent tracts of Neuropixel insertions. Neuropixels are dipped in Di-I resulting in a red fluorescent trace in the histology. Coronal slices at four different anterior-posterior positions are shown.

**Figure S2.**
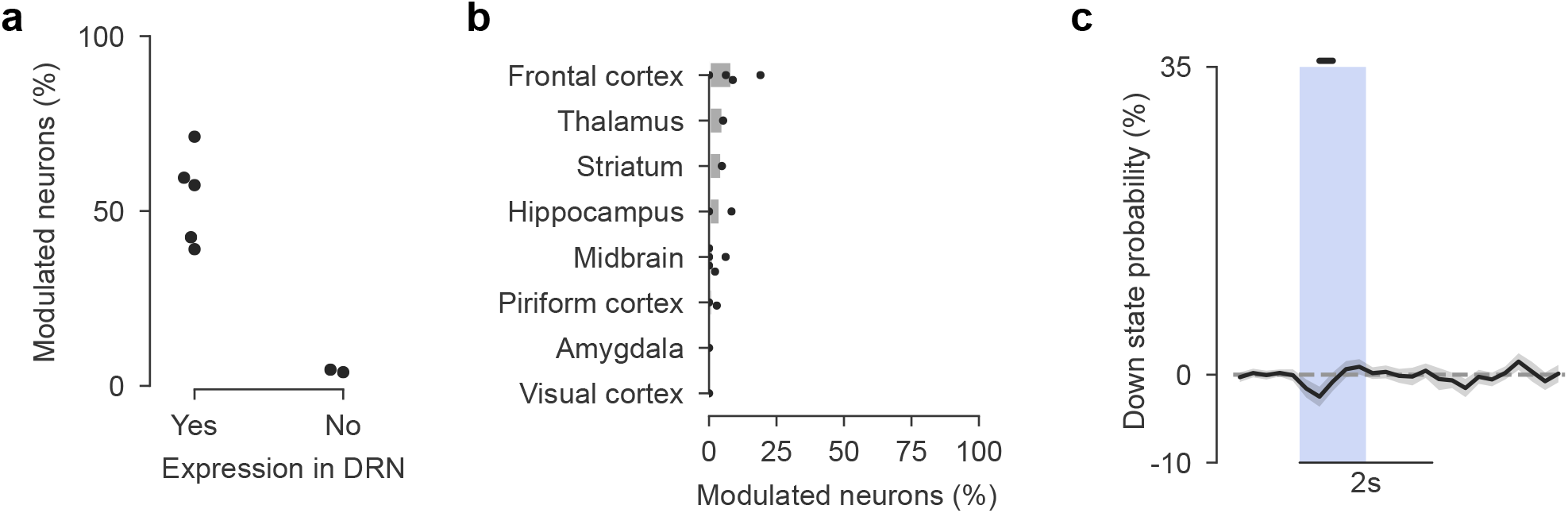
No light-evoked effect on neural activity in mice lacking channelrhodopsin expression. **(a)** The percentage of neurons which are significantly modulated by light delivery to the DRN. Each dot is a mouse, separated by whether that mouse expressed channelrhodopsin the DRN or not. **(b)** The percentage of light-modulated neurons in mice lacking channelrhodopsin expression, split out per brain region. Each dot is a recording session. **(c)** A very small decrease in DOWN state probability compared to baseline (−1 to 0s) was found in one time point during light delivery to the DRN (blue vertical bar). Data from all brain regions are pooled. Black horizontal bar above the plot indicates the significant time point (t-test versus 0, *p* = 0.046).

**Figure S3.**
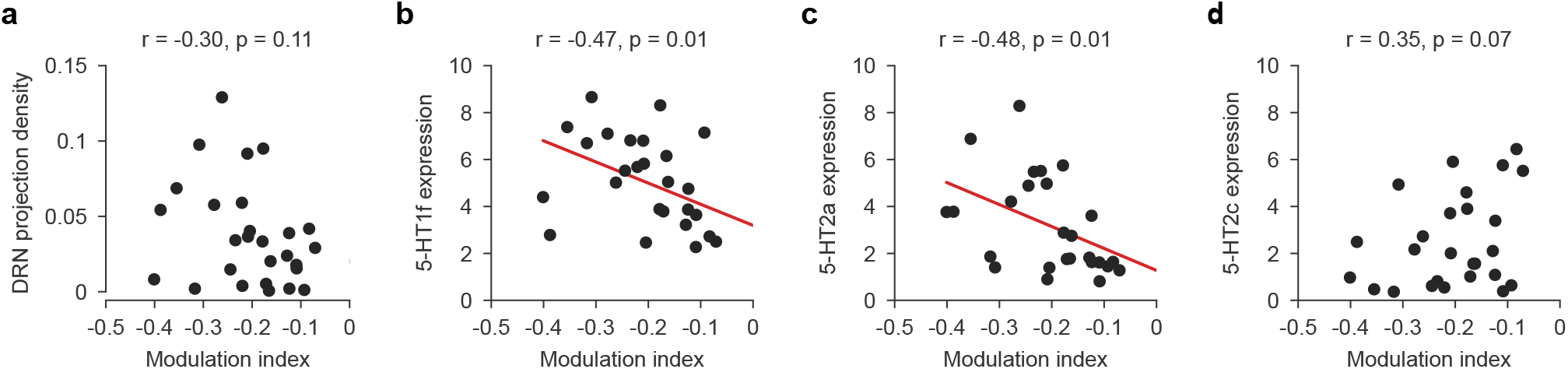
**(a)** No significant correlation between the projection strength from the dorsal raphe nucleus (DRN) with the magnitude of 5-HT induced suppression of neural activity. For each brain region, at a more granular delineation compared to the main figures, the projection density from the DRN was obtained from the Allen Brain Atlas. The modulation index of all neurons in each brain region was averaged. Each dot is a brain region. **(b)** There was a significant correlation for the expression strength of 5-HT1f receptors with 5-HT induced functional modulation at a region-by-region level. **(c)** Same as (b) for 5-HT2a receptor expression. **(d)** Same as (b) for 5-HT2c receptor expression. All stats done with a Pearson correlation.

**Figure S4.**
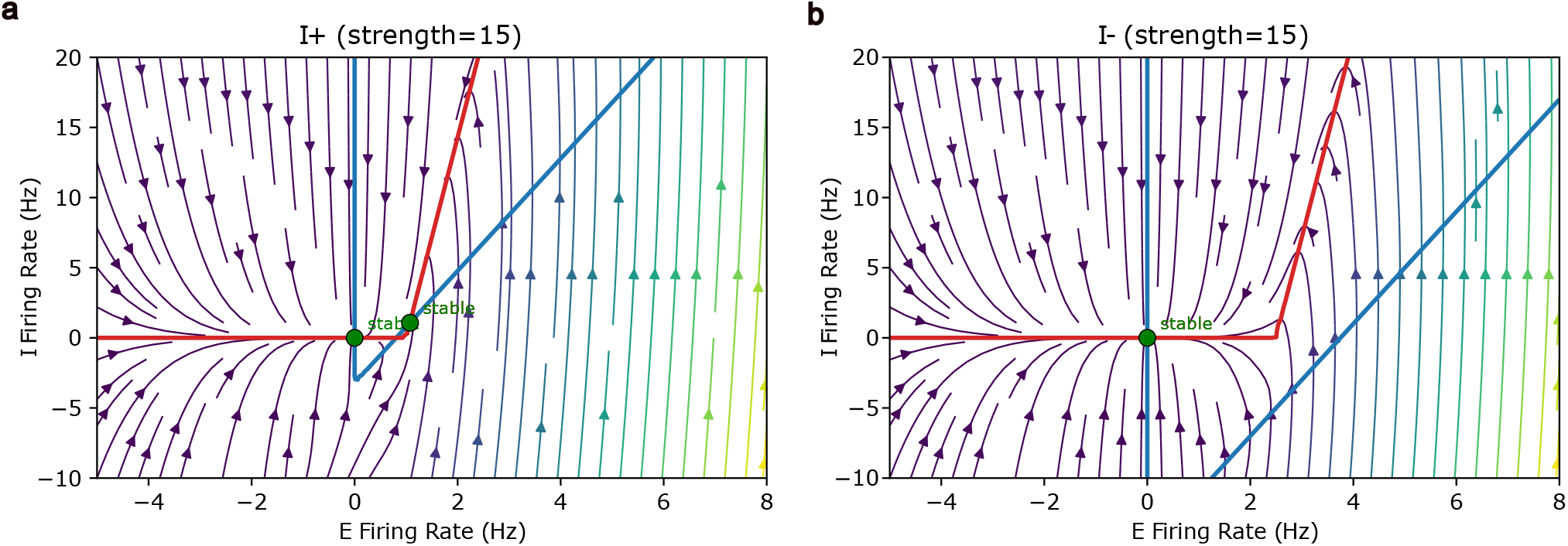
Nullclines of one isolated area containing one excitatory and one inihibitory population. **(a)** Excitatory external input (I+) to the inhibitory population with equal strength did not lead to the same DOWN state transition like inhibitory external input (I−) to the excitatory population. **(b)** Phase planes as in (a) for inhibitory input (I−).

**Figure S5.**
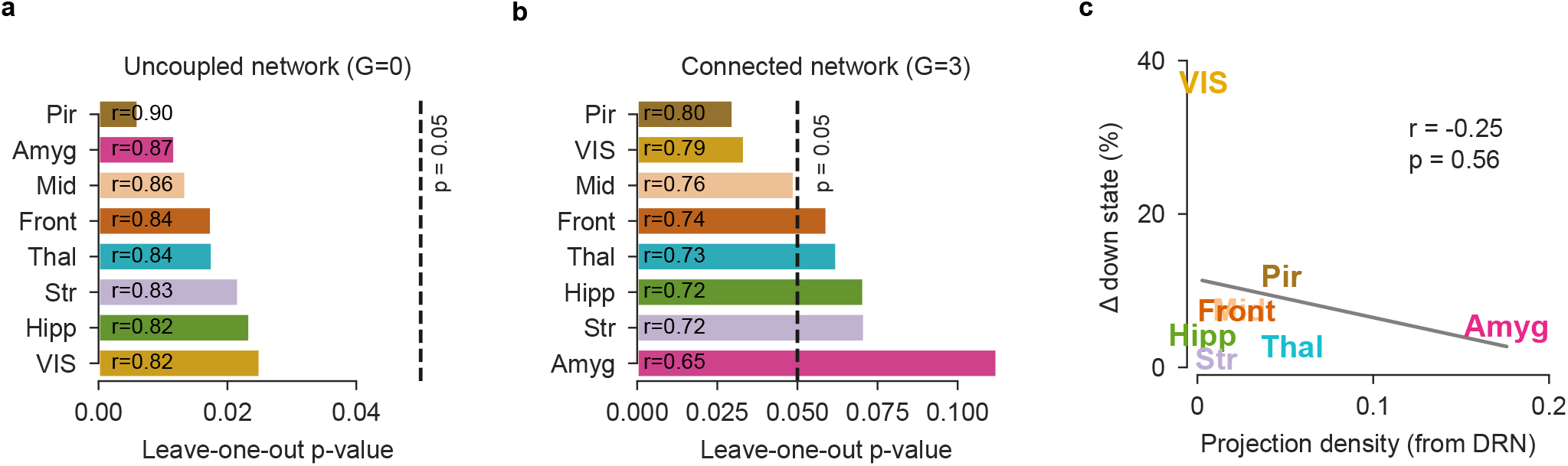
Robustness of DRN projection density correlations. **(a)** Leave-one-out analysis of the correlation between DRN projection density and DOWN state probability in the uncoupled network (*G*=0; corresponds to Fig. 3f, left). Each bar shows the Pearson correlation *p*-value obtained after excluding the labeled brain region from the correlation. The correlation remained significant regardless of which region was excluded (significance threshold *p*=0.05). **(b)** Same as (a) for the coupled network (*G*=3; Fig. 3f, right). Excluding several regions, most notably the amygdala, dominates the correlation, showing that this correlation. **(c)** The absolute change in DOWN state probability between the uncoupled (G=0) and coupled network (G=3) (Δ down state per region, as in Fig. 3h) plotted against DRN projection density. There was no significant correlation (Pearson correlation, *r*=-0.25, *p*=0.56).

## References

[1] P. Celada, M. Victoria Puig, and F. Artigas, Serotonin modulation of cortical neurons and networks, Frontiers in Integrative Neuroscience 7, 25 (2013), ISSN 16625145.

[2] M. Steriade, D. A. McCormick, and T. J. Sejnowski, Thalamocortical Oscillations in the Sleeping and Aroused Brain, Science 262, 679 (1993).

[3] M. Steriade, I. Timofeev, and F. Grenier, Natural Waking and Sleep States: A View From Inside Neocortical Neurons, Journal of Neurophysiology 85, 1969 (2001), ISSN 0022-3077, 1522-1598.

[4] G. T. Neske, The Slow Oscillation in Cortical and Thalamic Networks: Mechanisms and Functions, Frontiers in Neural Circuits 9, 88 (2015), ISSN 1662-5110.

[5] C. J. Wilson and P. M. Groves, Spontaneous firing patterns of identified spiny neurons in the rat neostriatum, Brain Research 220, 67 (1981), ISSN 0006-8993.

[6] M. V. Sanchez-Vives, M. Massimini, and M. Mattia, Shaping the Default Activity Pattern of the Cortical Network, Neuron 94, 993 (2017), ISSN 0896-6273.

[7] R. de Filippo, B. R. Rost, A. Stumpf, C. Cooper, J. J. Tukker, C. Harms, P. Beed, and D. Schmitz, Somatostatin interneurons activated by 5-HT2A receptor suppress slow oscillations in medial entorhinal cortex, eLife 10, e66960 (2021), ISSN 2050-084X.

[8] H. T. Hamada, Y. Abe, N. Takata, M. Taira, K. F. Tanaka, and K. Doya, Optogenetic activation of dorsal raphe serotonin neurons induces a brain-wide response in reward network, bioRxiv 2022.08.07.503074 (2022).

[9] J. Grandjean, A. Corcoba, M. C. Kahn, A. L. Upton, E. S. Deneris, E. Seifritz, F. Helmchen, E. O. Mann, M. Rudin, and B. J. Saab, A brain-wide functional map of the serotonergic responses to acute stress and fluoxetine, Nature Communications 2019 10:1 10, 1 (2019), ISSN 2041-1723.

[10] M. V. Puig, A. Watakabe, M. Ushimaru, T. Yamamori, and Y. Kawaguchi, Serotonin Modulates Fast-Spiking Interneuron and Synchronous Activity in the Rat Prefrontal Cortex through 5-HT1A and 5-HT2A Receptors, Journal of Neuroscience 30, 2211 (2010), ISSN 0270-6474.

[11] J. S. Montijn, K. Seignette, M. H. Howlett, J. L. Cazemier, M. Kamermans, C. N. Levelt, and J. A. Heimel, A parameter-free statistical test for neuronal responsiveness, eLife 10, e71969 (2021), ISSN 2050-084X.

[12] S. W. Oh, J. A. Harris, L. Ng, B. Winslow, N. Cain, S. Mihalas, Q. Wang, C. Lau, L. Kuan, A. M. Henry, et al., A mesoscale connectome of the mouse brain, Nature 508, 207 (2014), ISSN 1476-4687.

[13] J. E. Knox, K. D. Harris, N. Graddis, J. D. Whitesell, H. Zeng, J. A. Harris, E. Shea-Brown, and S. Mihalas, High-resolution data-driven model of the mouse connectome, Network Neuroscience 3, 217 (2018), ISSN 2472-1751.

[14] D. Jercog, A. Roxin, P. Barthó, A. Luczak, A. Compte, and J. De La Rocha, UP-DOWN cortical dynamics reflect state transitions in a bistable network, eLife 6 (2017), ISSN 2050084X.

[15] M. R. Joglekar, J. F. Mejias, G. R. Yang, and X. J. Wang, Inter-areal Balanced Amplification Enhances Signal Propagation in a Large-Scale Circuit Model of the Primate Cortex, Neuron 98, 222 (2018), ISSN 10974199.

[16] F. Deng, J. Wan, G. Li, H. Dong, X. Xia, Y. Wang, X. Li, C. Zhuang, Y. Zheng, L. Liu, et al., Improved green and red GRAB sensors for monitoring spatiotemporal serotonin release in vivo, Nature Methods 21, 692 (2024), ISSN 1548-7105.

[17] M. Torao-Angosto, A. Manasanch, M. Mattia, and M. V. Sanchez-Vives, Up and Down States During Slow Oscillations in Slow-Wave Sleep and Different Levels of Anesthesia, Frontiers in Systems Neuroscience 15 (2021), ISSN 1662-5137.

[18] J. Ren, D. Friedmann, J. Xiong, C. D. Liu, B. R. Ferguson, T. Weerakkody, K. E. DeLoach, C. Ran, A. Pun, Y. Sun, et al., Anatomically Defined and Functionally Distinct Dorsal Raphe Serotonin Sub-systems, Cell 175, 472 (2018), ISSN 0092-8674.

[19] G. T. Meijer, J. A. Catarino, L. Freitas-Silva, I. Laranjeira, I. B. Laboratory, and Z. F. Mainen, Serotonin drives choice-independent reconfiguration of distributed neural activity, bioRxiv 2025.08.01.668048 (2025), ISSN 2692-8205.

[20] I. B. Laboratory, K. Banga, J. Boussard, G. Chapius, M. Faulkner, K. D. Harris, J. Huntenburg, C. Hurwitz, H. Dong Lee, L. Paninski, et al., Spike sorting pipeline for the International Brain Laboratory (2022).

[21] S. Linderman, B. Antin, D. Zoltowski, and J. Glaser, SSM: Bayesian learning and inference for state space models, https://github.com/lindermanlab/ssm.

